# PTP1B phosphatase puts a brake on iPSC-derived neutrophil motility and antimicrobial function

**DOI:** 10.1101/2023.06.05.543775

**Authors:** Morgan A. Giese, David A. Bennin, Taylor J. Schoen, Ashley N. Peterson, Josh Brand, Ho Sun Jung, Huy Q. Dinh, Igor I. Slukvin, Anna Huttenlocher

## Abstract

Neutrophils are rapidly recruited to sites of infection and are critical for pathogen clearance. Neutropenic patients are at high risk for fungal and bacterial infections and can benefit from granulocyte transfusion therapy. Human induced pluripotent stem cells (iPSCs) could provide a robust source of neutrophil-like cells for infusion as they can be generated in large quantities and do not require a donor. However, dampened intracellular signaling limits their cellular activation and response. Here, we show that we can engineer iPSC-derived neutrophils (iNeutrophils) for enhanced motility and anti-microbial functions. Deletion of the PTP1B phosphatase increased iNeutrophil PI3K and ERK signaling and was associated with increased F-actin polymerization, cell migration and phagocytosis. PTP1B deletion also increased production of inflammatory cytokines, including the neutrophil chemoattractant IL-8. Furthermore, PTP1B-KO iNeutrophils displayed a highly activated morphology and were more responsive to the fungal pathogen *Aspergillus fumigatus*. KO iNeutrophils efficiently migrated to and swarmed hyphae resulting in inhibition of fungal growth. Taken together, deletion of the PTP1B phosphatase removes the “brakes” on iPSC-derived intracellular signaling and neutrophil function.

**Key points:** Deletion of PTP1B increases iPSC-derived neutrophil intracellular signaling to improve motility and phagocytosis.

Deletion of PTP1B enhances iPSC-derived neutrophil swarming response and ability to inhibit fungal growth.

## Introduction

Neutrophils are highly motile cells and as first responders of innate immunity are critical for clearing bacterial and fungal infections. They exhibit an arsenal of anti-microbial functions, including production of reactive oxygen species (ROS), release of neutrophil extracellular traps (NETs), and phagocytosis of pathogens. Patients who have received bone marrow transplant or chemotherapy can become neutropenic, leading to higher rates of mortality due to increased susceptibility to bacterial and fungal infections ^1^. Alongside administration of antifungals or antibiotics, granulocyte transfusion therapy (GTX) can be utilized to improve patient outcomes. However, success of GTX has suffered from donor yield, as well as the short lifespan and functional capacity of primary human neutrophils ex vivo ^2^. Thus, there is need for identification of alternative sources of infusible neutrophils.

Induced pluripotent stem cells (iPSC)-derived neutrophils (iNeutrophils) have emerged as a promising option for treating neutropenic patients as no donor is required, the cells are genetically malleable, and can be generated in large numbers. iNeutrophils have been shown to migrate effectively and exert many neutrophil antimicrobial functions ^3-6^. However, their functional capacity is often lower than primary human neutrophils ^3-6^. iNeutrophils were shown to have dampened activation of intracellular signaling pathways, such as phospho-AKT, that limit functions including migration, phagocytosis and NETosis ^6^. Thus, inhibitory signaling pathways may prevent full activation and response. Methods to activate intracellular signaling pathways may improve iNeutrophil function for future use as a clinical therapy for neutropenic patients. Furthermore, iNeutrophils can be a valuable tool to study the molecular signaling pathways that regulate neutrophil migration and antimicrobial function.

Here, we show that we can engineer iPSC-derived neutrophils to enhance cellular activation, motility and antimicrobial response. Protein Tyrosine Phosphatases such as PTP1B negatively regulate many signaling pathways involved in neutrophil function ^7^. Deletion of the PTP1B phosphatase increased iNeutrophil PI3K and ERK signaling leading to increased F-actin polymerization, cell migration and phagocytosis. PTP1B-KO iNeutrophils produced higher levels of inflammatory chemokines, including the neutrophil chemoattractant IL-8. Furthermore, PTP1B-KO iNeutrophils displayed a highly activated morphology in the presence of the fungal pathogen *Aspergillus fumigatus*, efficiently migrated to and swarmed hyphae, and limited fungal growth. Thus, deletion of the PTP1B phosphatase removes the “brakes” on iPSC-derived intracellular signaling and neutrophil function.

## Methods

### Stem Cell Culture and Neutrophil Differentiation

Neutrophils were differentiated from the bone marrow-derived IISH2i-BM9 cell line (WiCell) ^8^. hiPSCs were cultured on Matrigel-coated plates in mTeSR-Plus medium. To induce hemogenic endothelium, hiPSCs are transfected with *ETV2* mmRNA. One day following transfection, media was changed to StemLineII with VEGF-165 and FGF to induce differentiation into hemogenic endothelial cells. After two days, media was changed to StemLineII supplemented with FGF2, GM-CSF, and UM171 to generate common myeloid progenitors (CMPs). On days 8-10, floating cells were gently harvested and cultured in StemSpan SFEM II supplemented with GlutaMAX, ExCyte, G-CSF and Am580 to promote terminal neutrophil differentiation. Floating neutrophils were harvested after another 8-10 days.

### Generation of PTP1B^-/-^ BM9iPSCs

Cells were nucleofected with Cas9 protein and two guide RNAs targeting Exon 3 of *PTPN1*. Individual colonies were picked and expanded. To confirm biallelic mutation, clones were screened by PCR for the acquisition of a 67 bp deletion. Loss of protein expression was verified by western blot.

### Human Neutrophil Isolation

Human blood was obtained from volunteering donors with informed written consent through a protocol that was approved by the Internal Review Board of the University of Wisconsin-Madison. Neutrophils were isolated using MACSxpress negative antibody selection kit and purified with the MACSxpress erythrocyte depletion kit.

### Flow Cytometry

Neutrophils were stained **(Supplementary Table 1)**, fixed and acquired on an Aurora Cytometer. Myeloid cells were identified by CD11b+ expression and neutrophils were identified by CD15+ or CD15+CD16+ expression. Monocytes were identified as CD14+. Data were analyzed using FlowJo Software (v10.8.1).

#### Flow cytometry analysis in R using Cytofworkflow

Exported .fcs files were analyzed with *prepData* function from Cytofworkflow (v1.14.0) pipeline ^9^. Data transformation was calculated for all fluorescent parameter using an *arcsinh* normalization and a cofactor of 3000. Transformed data were plotted using a smoothed density function using ggplot2 to show proportion of cell types over the range of expression values.

### Presto Blue Viability

Cells were plated and incubated at 37°C. For each timepoint, Presto Blue HS was added and incubated for 30 minutes before reading fluorescence at 560/590 nm in a microplate reader.

### Western Blot Cell Signaling

Cells were stimulated with 1uM fMLP for 3 minutes then collected in lysis buffer with protease and phosphatase inhibitors. Cells were sonicated and clarified by centrifugation. Protein concentrations were determined and immunoblotting of cell lysates was performed **(Supplementary Table 2)** and blots were imaged with an infrared imaging system.

### Chemotaxis

Chemotaxis was assessed using a microfluidic device ^10^. Polydimethylsiloxane (PDMS) devices were adhered to glass coverslips and coated with fibrinogen. Calcein stained cells were added before addition of fMLP. Cells were imaged every 30 seconds for 45–90 min on an inverted fluorescent microscope with a 10× objective and an automated stage. Cell tracking analysis was done using JEX software ^11^.

### Immunofluorescent imaging

Cells were stimulated in an fMLP bath for 30 minutes at 37°C, then fixed, permeabilized, and incubated with Rhodamine-phalloidin overnight at 4°C. Coverslips were counterstained with Hoechst 33342 and imaged on an upright Laser Scanning Confocal Microscope equipped with a motorized stage. Images were acquired with a 60× oil, NA 1.40 objective and processed in ZenBlue software. Maximum intensity projections were generated and integrated density of the actin channel was quantified using ImageJ.

### Phagocytosis

pHrodo™ Green *E. coli* BioParticles were opsonized with 30% pooled human serum and incubated with iNeutrophils (100:1 MOI) for 1 hour at 37°C. Reaction was stopped by addition of ice cold PBS. Cells were stained for flow cytometry **(Supplementary Table 1**) on ice, then fixed before analysis using an Aurora Cytometer.

### NETosis and Intracellular ROS

For NETosis, cells were stimulated with PMA and incubated for 4 hours at 37°C before endpoint addition of Sytox Green. For ROS assay, DHR123 probe was added before PMA and plate was read every 15 minutes after stimulation. NETosis (500/528nm) and ROS (485/535nm) fluorescence was quantified using a microplate reader.

### Real Time qPCR

Cells were stimulated with 200ng/mL *E. coli* LPS for 2 hours at 37°C, then collected in Trizol for RNA isolation. cDNA was synthesized and used as the template for quantitative PCR (qPCR) **(Supplementary Table 3)**. Data were normalized to *ef1a* within each sample using the ΔΔCq method ^12^. Fold-change represents the change in cytokine expression over the unstimulated WT sample.

### Inflammatory Cytokine and LTB4 ELISA

For inflammatory cytokines IL-8, IL-1beta, IL-6 and TNFa, iNeutrophils were stimulated with 200ng/mL *E. coli* LPS or 10ug/mL Zymosan for 4 hours at 37°C. For LTB4, iNeutrophils were stimulated with 1uM fMLP for 5 or 30 minutes at 37°C. Supernatants were collected, aliquoted and frozen at -80°C until quantification by ELISA.

### Fungal co-culture imaging

*Aspergillus fumigatus* (CEA10) was grown as previously described ^13^. For imaging experiments, *A. fumigatus* were incubated until germling stage (37°C for 8 hours). GMM media was removed and replaced with neutrophil suspension (MOI 150:1). Images were taken every 3 minutes on an inverted fluorescent microscope with a 20× objective and an automated stage at 37°C with 5% CO_2_. Videos were compiled using ImageJ software.

#### Image Analysis

iNeutrophil *circularity* and cluster size was quantified by outlining individual cells or clusters using the Polygon selection tool (ImageJ). A cluster was identified as a tightly formed group of at least 5 iNeutrophils attached to *A. fumigatus*. Hyphal length was measured using the segmented line feature (ImageJ).

### Statistical Analysis

All experiments and statistical analyses represent at least three independent replicates. Analysis of chemotactic index and cell velocity was performed using unpaired Student’s t-test. Analysis of receptor expression by flow cytometry, phagocytosis, qPCR gene expression and ELISA was performed using paired Student’s t-test. Analysis of Presto Blue viability, ROS production, and percent of clustered germlings was performed using simple linear regression and the p value was calculated by comparing the slope of each line. Analysis of cluster size over time was performed by calculating the area under the curve (AUC) for each replicate and then determining the p value using a paired Student’s t-test. Analysis of NETosis and phospho-signaling was performed using One Sample t-test. Above analyses were conducted using GraphPad Prism (v9). Integrated Density of F actin, circularity, and hyphal length analysis represent least-squared adjusted means±standard error of the mean (LSmeans±s.e.m.) and were compared using ANOVA with Tukey’s multiple comparisons (RStudio).

## Results

### Generation of PTP1B-KO iPSC-derived neutrophils

To increase activation of intracellular signaling pathways in iPSC-derived neutrophils, we deleted the protein tyrosine phosphatase 1B (PTP1B), encoded by *PTPN1*, using CRISPR/Cas9 mediated gene mutation at the iPSC-stage (Figure 1A). We generated clonal cell lines with biallelic deletion of *PTPN1* (Supplementary Figure 1A), and then differentiated these cells into human iPSC-derived neutrophils using serum-and feeder-free conditions, following published protocols ^5,8^ (Figure 1B). Loss of PTP1B protein expression was confirmed both at the stem cell stage and after neutrophil differentiation by western blot (Supplementary Figure 1B, Figure 1C). Morphological characteristics of differentiated neutrophils were examined by cytospin, confirming the presence of hyper-segmented nuclei (Figure 1D).

**Figure 1.**
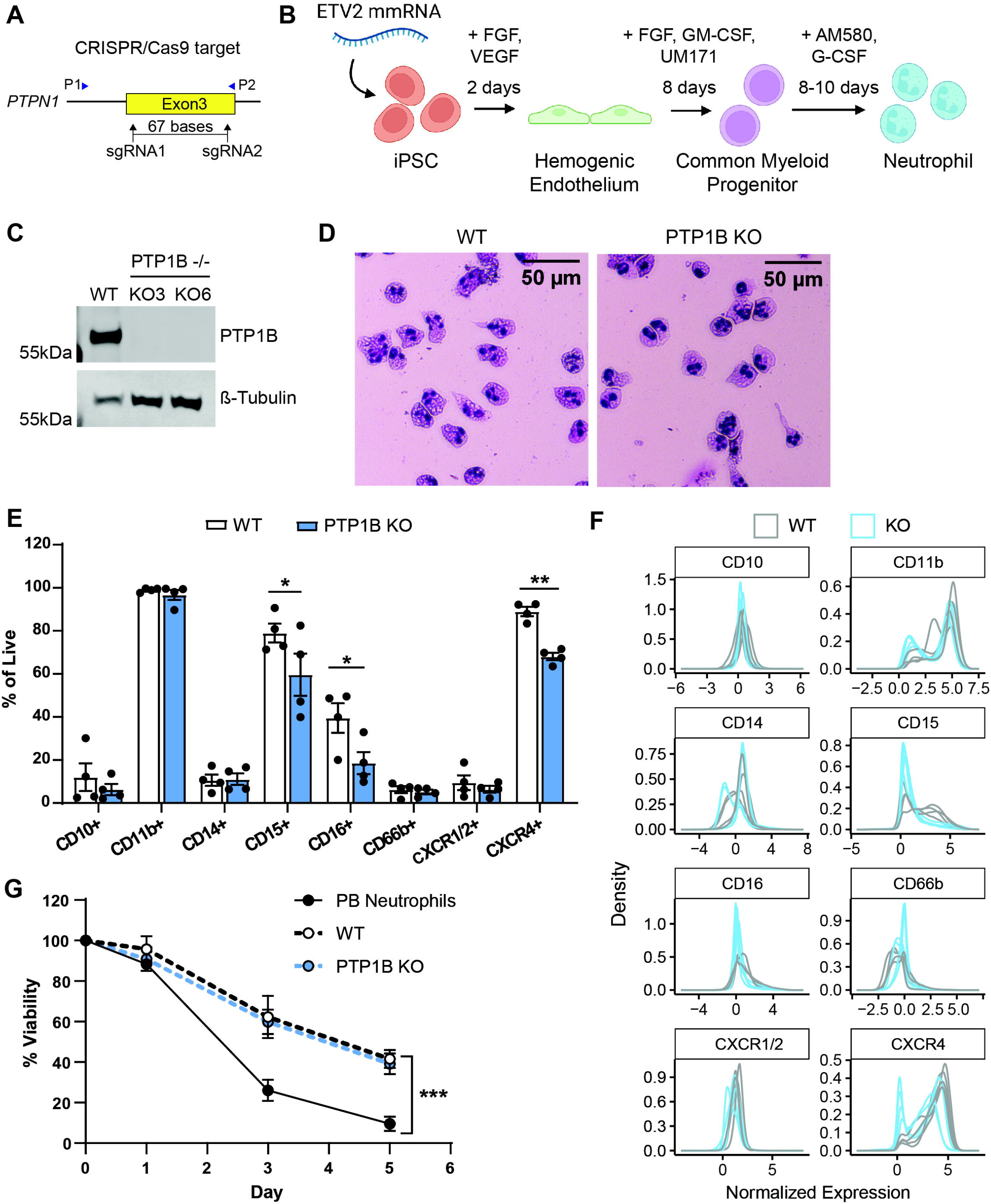
Generation of PTP1B-KO iPSC-derived neutrophils. (A) Diagram illustrates sgRNAs targeting exon 3 of *PTPN1* for CRISPR/Cas9 mediated deletion of a 67bp region at the stem cell stage. (B) Timeline for neutrophil differentiation from bone marrow derived iPSCs. (C) Western blot confirmation of PTP1B CRISPR/Cas9 mediated deletion in differentiated neutrophils. (D) Representative cytospins showing morphological confirmation of neutrophil differentiation. (E) Flow cytometry staining of differentiated neutrophils and myeloid cells. Cells were gated on live cells. (F) Histogram plots of normalized flow cytometry surface receptor expression data in (E). Data were normalized individually to each fluorescent marker and presents the range of expression. (G) Cell viability of iPSC-derived neutrophils compared to human peripheral blood (PB) neutrophils. Diagram (B) was created with BioRender.com. Experiments were conducted at least three times, or as indicated on the plot. Dots in (E) represent independent replicates and dots in (G) represent the average of all replicates. Means ± SEM are shown. *p* values were calculated by paired Student’s t-test (E) or by simple linear regression (G). *, p < 0.05; **, p < 0.01

Further validation of neutrophil differentiation was completed by staining for neutrophil surface receptors. As previously reported, almost all iNeutrophils expressed the common myeloid marker CD11b, but less than 20% expressed the canonical human neutrophil marker CD66b or mature neutrophil marker CD10. However, the majority of iNeutrophils expressed primary blood and mature neutrophil markers CD15 and CD16 (Figure 1E, F) ^3,5^. While deletion of PTP1B resulted in a lower proportion of fully mature CD16+ neutrophils, the majority of PTP1B-KO cells still expressed CD15 (Figure 1E). PTP1B has been shown to modify murine myelopoiesis by negatively regulating monocyte differentiation ^14^, however, we did not find a significant increase in CD14+ monocytes upon PTP1B deletion in human iPSCs (Figure 1E). Our data indicate that deletion of PTP1B at the stem cell stage still allows for neutrophil differentiation.

A limitation of primary human peripheral blood (PB) neutrophils is their short life span ex vivo. To determine if the lifespan of iNeutrophils is longer than that of PB neutrophils, we evaluated the longevity of these cells in culture. PB neutrophils exhibited less than 30% viability at 3 days, whereas iNeutrophils were 50% viable after 5 days, with no difference between WT and PTP1B-KO cells (Figure 1G). The lower maturation level of our iNeutrophils may explain the longer lifespan of these cells compared to fully matured PB neutrophils.

### Deletion of PTP1B promotes intracellular signaling, motility and actin polymerization

Previously we and others have reported that PTP1B regulates the actin cytoskeleton and inhibits migration of cancer cells ^15,16^. To determine if PTP1B also regulates signaling and motility of human iNeutrophils, we first quantified phospho-signaling induced by the bacterial formylated peptide fMLP. PTP1B negatively regulates MAPK and PI3K signaling pathways ^17^. PTP1B-KO iNeutrophils showed increased phosphorylation of ERK1/2, AKT, and HS1 compared to WT iNeutrophils (Figure 2A, B). HS1 is an actin binding protein that regulates actin dynamics during cell migration via its interaction with the Arp2/3 complex. Enhanced phosphorylation of HS1 is correlated with efficient directed neutrophil migration ^18^. Using a previously published microfluidic device ^19^, we conducted short term live imaging of neutrophil migration in response to the chemoattractant fMLP. PTP1B-KO iNeutrophils displayed enhanced motility with higher chemotactic index and velocity compared to WT cells (Figure 2C).

**Figure 2.**
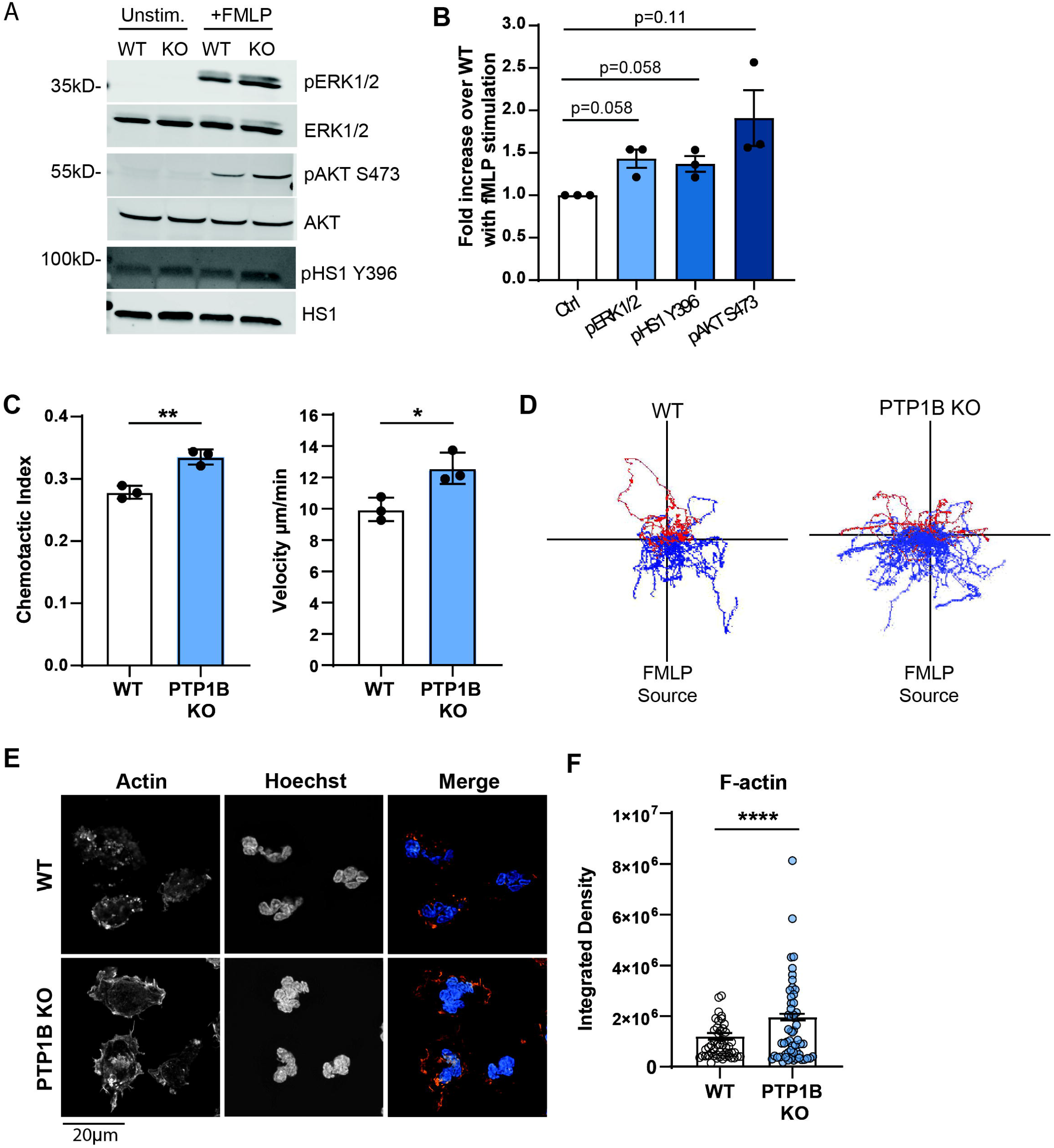
Deletion of PTP1B promotes intracellular signaling, motility and actin polymerization. (A) Representative western blots and quantification (B) of ERK, HS1, AKT phospho-signaling after stimulation with 1uM fMLP for 3 minutes. (C) iNeutrophil chemotactic index and mean velocity in response to an fMLP gradient over 45 minutes of imaging. (D) Representative track plots of cells migrating in response to an fMLP gradient. Blue tracks indicate cells that traveled towards the fMLP source, whereas red tracks indicate cells that moved away. (E) Representative immunofluorescence images of F-actin staining after 100nM fMLP stim. (F) Quantified integrated density of F-actin staining. Dots represent individual cells. WT n=50, KO n=56. Scale bar represents 20μm. Experiments were conducted at least three times, or as indicated on the plot. Dots in (B, C) represent independent replicates. Bars in (B, C) represent means ± SEM. Bars in (F) represent LSmeans ± SEM. *p* values were calculated by One-sample t-test (B), unpaired Student’s t-test (C) or ANOVA with Tukey’s multiple comparisons (F). *, p < 0.05; **, p < 0.01; ****, p < 0.0001.

Representative cell tracks show increased directed migration of PTP1B-KO iNeutrophils (Figure 2D). During migration, actin is polymerized at the leading edge to drive pseudopod formation ^20^. As elevated p-HS1 may increase actin polymerization to promote PTP1B-KO iNeutrophils chemotaxis, we next imaged F-actin polymerization in iNeutrophils after stimulation in an fMLP bath. PTP1B-KO cells displayed increased density of total F-actin compared to WT cells by immunofluorescent staining (Figure 2E, F). Thus, deletion of the PTP1B phosphatase increases intracellular ERK and PI3K signaling and the motility of iNeutrophils.

### Deletion of PTP1B improves neutrophil phagocytosis but decreases production of ROS and NETs

Increased intracellular phospho-signaling may promote neutrophil antimicrobial functions, including the ability to phagocytose microbes ^7^. Furthermore, phagocytosis is mediated by actin contraction to form the phagosome ^21^, and thus, we hypothesized this function may be enhanced in PTP1B-KO iNeutrophils. We quantified phagocytosis of *E. coli* coated beads by flow cytometry and found a significant increase in phagocytosis by CD15+ PTP1B-KO iNeutrophils (Figure 3A). This effect was heightened when gating on CD15+ CD16+ mature neutrophils with 80% of PTP1B-KO iNeutrophils phagocytosing and acidifying *E. coli* coated beads, whereas less than 50% of WT cells had after 1 hour.

**Figure 3.**
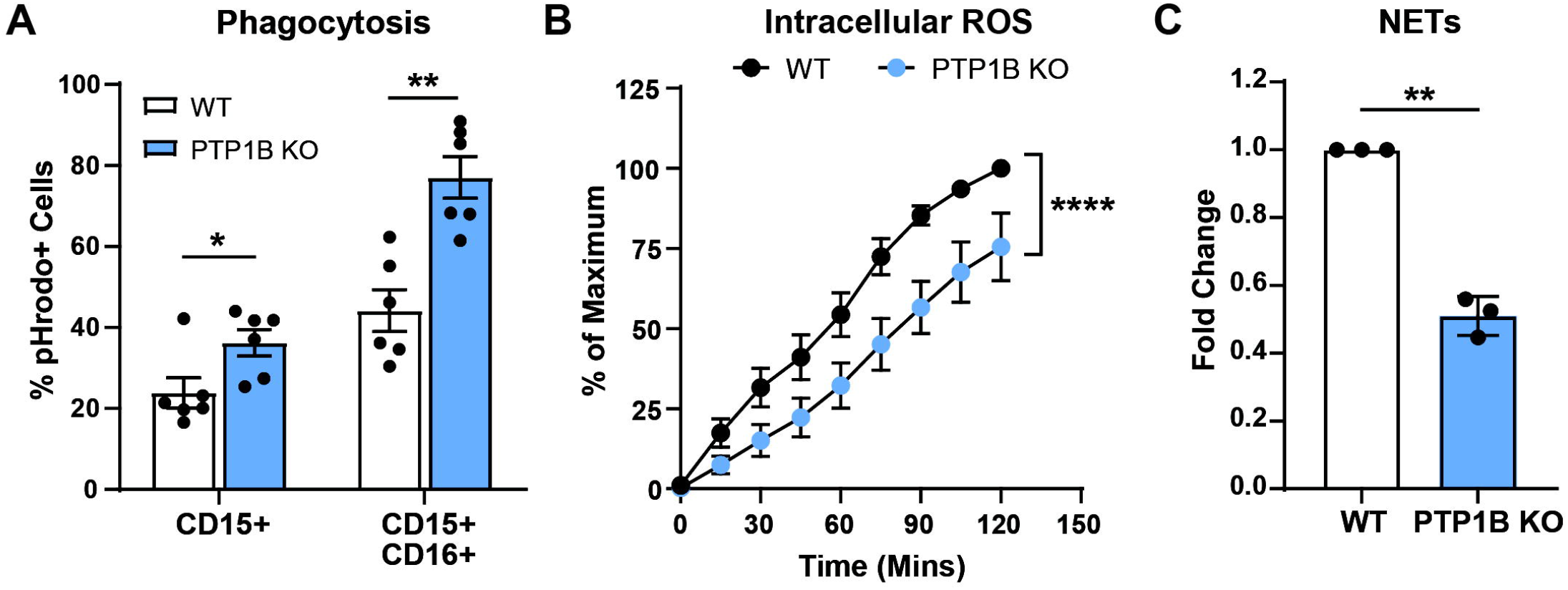
Deletion of PTP1B improves neutrophil phagocytosis but decreases production of ROS and NETs. (A) iNeutrophil phagocytosis of acidified pHrodo *E. coli* beads quantified by flow cytometry. Percent pHrodo+ cells of CD11b+CD15+ neutrophils or CD11b+CD15+CD16+ mature neutrophils. (B) Quantification of iNeutrophil intracellular ROS production over time using DHR123 peroxynitrite indicator following stimulation with 50ng/mL PMA. (C) NETosis quantified with Sytox Green DNA indicator after 4 hour stimulation with 100ng/mL PMA. Experiments were conducted at least three times, or as indicated on the plot. Dots in (A, C) represent independent replicates and dots in (B) represent the average of all replicates. Means ± SEM are shown. *p* values were calculated by paired Student’s t-test (A), simple linear regression (B), or One-sample t-test (C). *, p < 0.05; **, p < 0.001; ****, p < 0.0001.

Following phagocytosis, neutrophils can kill pathogens by production of intracellular reactive oxygen species (ROS). While PTP1B-KO iNeutrophils produced lower levels of ROS than WT cells, PTP1B-KO iNeutrophils were still capable of producing ROS at high levels upon PMA stimulation (Figure 3B). It is likely the PTP1B-KO iNeutrophils produce levels similar to that of human PB neutrophils (Supplementary Figure 2A). Another antimicrobial mechanism is the release of neutrophil extracellular traps (NETs). Upon PMA stimulation, we found that PTP1B-KO iNeutrophils make fewer NETs (Figure 2E). ROS production is necessary for NET release ^22^, thus, the decreased production of NETs correlates with the decrease in ROS we observed in PTP1B-KO iNeutrophils.

### Deletion of PTP1B increases inflammatory cytokine production

Upon migrating to sites of infection, neutrophils can produce inflammatory cytokines to further recruit and promote the innate and adaptive immune response. IL-8 is a strong neutrophil chemoattractant and activating factor. *CXCL8* transcripts were increased both basally and upon LPS stimulation in PTP1B-KO iNeutrophils (Supplemental Figure 3A), indicating that PTP1B negatively regulates IL-8 expression. We confirmed increased production at the protein level with both LPS and Zymosan stimulation (Figure 4A). Increased production of IL-8 may contribute to enhanced iNeutrophil activation and chemotactic response during bacterial or fungal infection. We also found differential production of the inflammatory cytokines TNFα, IL1β and IL-6. PTP1B-KO iNeutrophils showed comparable levels of *IL1B* at the basal level (Supp. Figure 3B), but significantly increased protein secretion upon stimulation (Figure 4B). Additionally, PTP1B-KO iNeutrophils showed elevated secretion of IL-6 and TNFα after stimulation (Supplemental Figure 3C, D, Figure 4C, D). Taken together these findings suggest that PTP1B negatively regulates inflammatory cytokine production.

**Figure 4.**
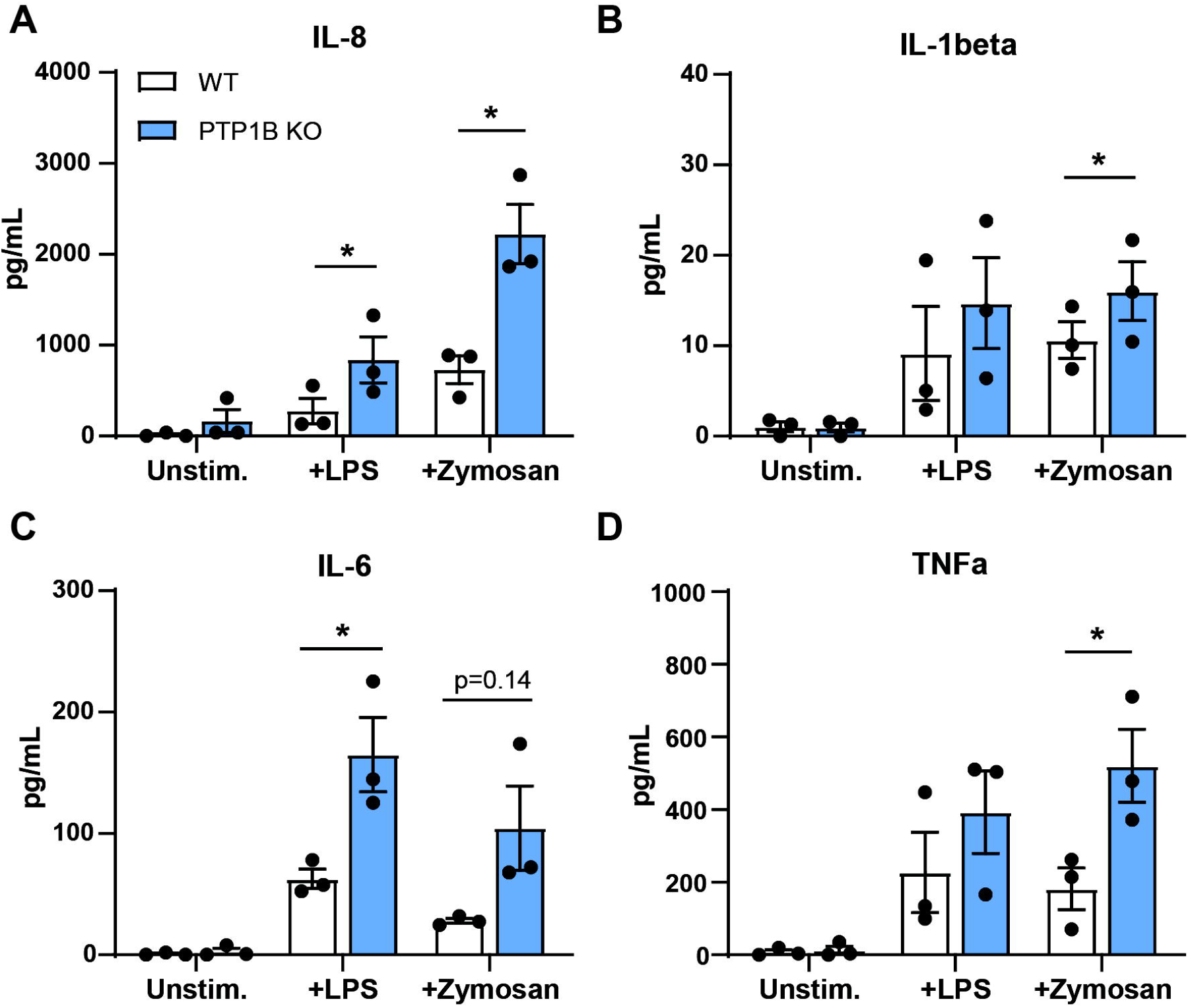
Deletion of PTP1B increases inflammatory cytokine production. (A-D) Inflammatory cytokines produced by unstimulated iNeutrophils or after stimulation with 200ng/mL LPS or 10ug/mL Zymosan for 4 hours. Experiments were conducted three times. Dots represent independent replicates. Means ± SEM are shown. *p* values were calculated by paired Student’s t-test (A-D). *, p < 0.05.

To determine if increased cytokine production by PTP1B-KO iNeutrophils is due to altered expression of pathogen recognition receptors (PRRs), rather than increased intracellular signaling, we quantified expression of Dectin-1, TLR2, and TLR4 on WT and KO iNeutrophils. Zymosan is recognized by Dectin-1 and TLR2, whereas LPS is primarily recognized by TLR4 ^23^. We found similar levels of expression of these receptors on WT and KO cells, thus the increase in inflammatory cytokine production is not due to differences in PRR expression (Supplementary Figure 3E). Our findings show that PTP1B-KO cells are more responsive to microbial induced production of inflammatory cytokines compared to WT iNeutrophils.

### Deletion of PTP1B increases iNeutrophil swarming and inhibition of fungal growth

After evaluating the effect of PTP1B deletion on specific neutrophil functions, we next wanted to determine if PTP1B expression affects iNeutrophil response to the live fungal pathogen *Aspergillus fumigatus*. We co-cultured iNeutrophils with *A. fumigatus* at the germling stage and then live imaged iNeutrophil-fungal interactions over the course of 8 hours (Figure 5A). We noticed a stark difference in the morphology of WT vs PTP1B-KO iNeutrophils in the presence of *A. fumigatus*. During co-incubation, KO iNeutrophils displayed an elongated cell shape indicative of cell activation (Figure 5B). Specifically, PTP1B-KO iNeutrophils showed significantly decreased circularity in the presence of *A. fumigatus* (Figure 5C). In contrast, the majority of WT cells remained maintained a round morphology, even after 4 hours (Figure 5B, C). Limited activation of WT iNeutrophils in response to *A. fumigatus* is likely not due to decreased recognition, as we saw no difference in Dectin-1 expression (Supplementary Figure 3E) but may be due to dampened signaling mediated by PTP1B.

**Figure 5.**
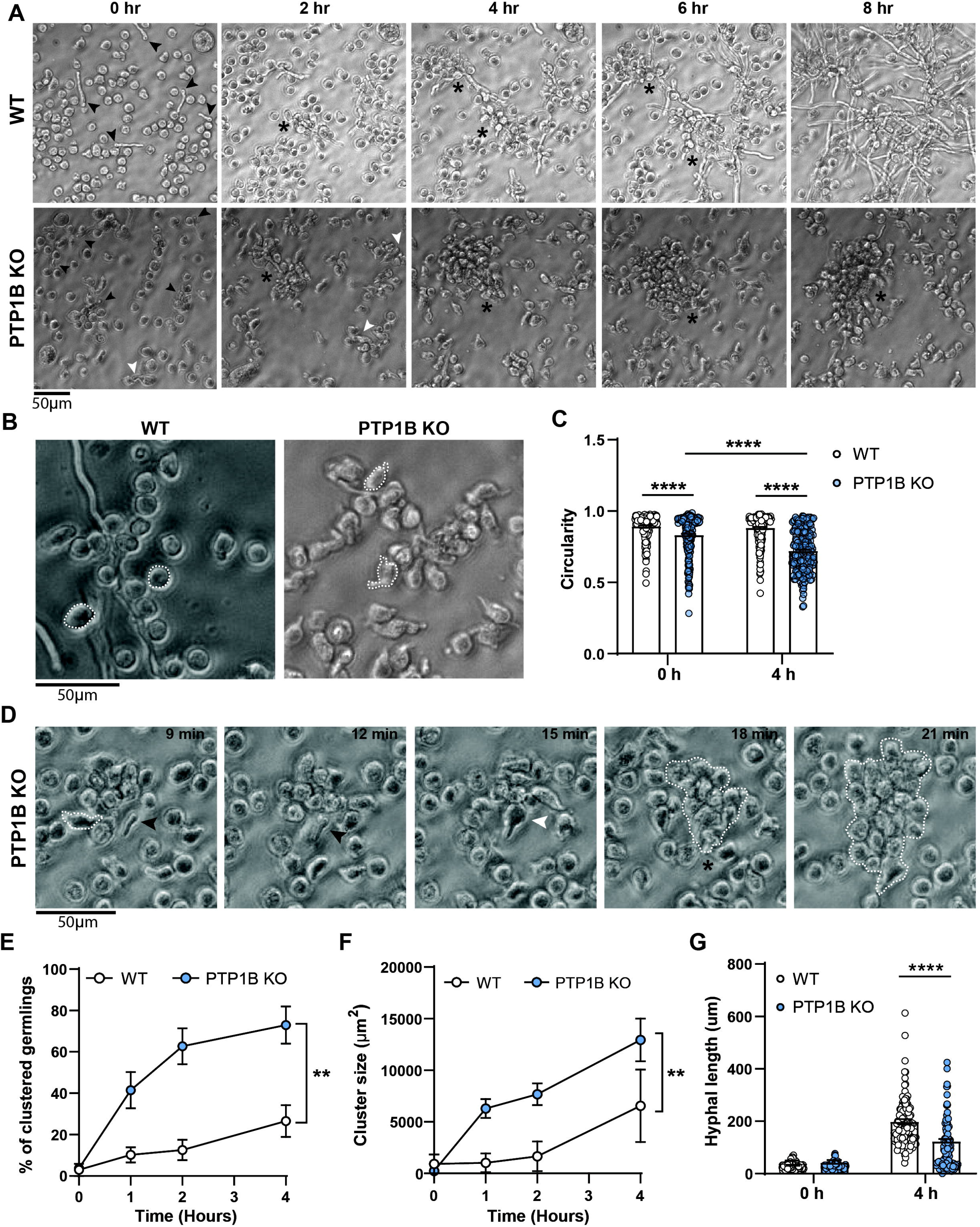
Deletion of PTP1B increases iNeutrophil swarming and inhibition of fungal growth. (A) Representative bright-field images of *A. fumigatus* co-culture with WT or PTP1B-KO iNeutrophils over the course of 8 hours. Black arrows indicate germlings. White arrows indicate a phagocytosed germling. Asterisks indicate iNeutrophil swarming and cluster formation. (B) Higher magnification bright-field images of WT or PTP1B-KO iNeutrophil cell morphology and interaction with *A. fumigatus* hyphae. Individual cells are outlined. (C) Quantification of iNeutrophil cell shape (circularity) at 0 and 4 hours of co-incubation. Dots represent individual cells. WT n=414, KO n=396. (D) Representative time lapse images of PTP1B-KO iNeutrophils phagocytosing and clustering around *A. fumigatus* hyphae. Black arrows indicate germlings. White arrows indicate a phagocytosed germling. Asterisks indicate iNeutrophil swarming and cluster formation. Clusters are outlined. (E) Quantification of percent of *A. fumigatus* germlings surrounded by iNeutrophil clusters at 0, 1, 2, and 4 hours of co-incubation. (F) Quantification of iNeutrophil cluster size at 0, 1, 2, and 4 hours of co-incubation. (G) Quantification of hyphal length at 0 and 4 hours of co-culture with WT or PTP1B-KO iNeutrophils. Dots represent individual germlings. WT n=138, KO n=110. Scale bars represent 50μm. Experiments were conducted at least three times, or as indicated on the plot. Bars in (C,G) represent LSmeans ± SEM. Dots in (E, F) represent the mean± SEM. *p* values were calculated by ANOVA with Tukey’s multiple comparisons (C, G), linear regression (E), or paired Student’s t-test (F) on the calculated area under the curve (AUC) for each replicate. **, p < 0.01; ****, p < 0.0001.

We next analyzed iNeutrophil motility in response to *A. fumigatus* fungal growth. We found that PTP1B-KO iNeutrophils were highly migratory and rapidly recruited and swarmed *A. fumigatus* (Video S1). In contrast, WT iNeutrophils showed a delayed response with fewer cells migrating and physically interacting with germlings or hyphae (Video S2, S3). Furthermore, we observed many instances of PTP1B-KO iNeutrophils phagocytosing germlings that led to enhanced neutrophil recruitment and swarming (Figure 5D and Video S4, S5). PTP1B-KO iNeutrophils formed tight clusters around *A. fumigatus* significantly faster and larger in size than WT cells (Figure 5E, F). Neutrophil swarming is mediated by release of LTB4 ^24^, therefore, increased swarming by PTP1B-KO iNeutrophils could indicate increased production of LTB4 or expression of its receptor BLT1R. We found that almost 100% of WT and KO iNeutrophils expressed BLT1R but found no difference in the level of expression between genotypes (Supplementary Figure 4A, B). We quantified LTB4 release upon fMLP stimulation and saw no significant difference between WT and KO cells (Supplementary Figure 4C). Thus, swarming by PTP1B-KO iNeutrophils is likely due to enhanced cell activation that leads to migration and physical interaction with *A. fumigatus*.

Lastly, we wanted to determine if heightened PTP1B-KO iNeutrophil recruitment and swarming of *A. fumigatus* limited fungal growth. We quantified the hyphal length for each germling at 0 and 4 hours of co-incubation. After 4 hours, hyphal growth was significantly decreased when cultured with PTP1B-KO neutrophils, compared to WT cells (Figure 5G). Taken together, our findings demonstrate that PTP1B-KO iNeutrophils are more responsive to *A. fumigatus* resulting in enhanced recruitment, swarming and inhibition of fungal growth.

## Discussion

iPSC-derived neutrophils can exert many classic primary neutrophil functions. In vivo mouse studies using infusible iNeutrophils show promise for treating diseases ranging from bacterial infection to cancer ^4,6,25-27^. However, many of these iNeutrophils display inhibited or lower functional capacity compared to primary human neutrophils ^3-6^. Therefore, there is a need to improve our understanding of the molecular signaling pathways that regulate iNeutrophil function to enhance their use as a clinical therapy. iNeutrophils can be genetically modified and thus are a valuable tool for dissecting pathways regulating cell migration and antimicrobial response. In this study, we genetically manipulated iNeutrophils and found that deletion of the phosphatase PTP1B increases cellular activation and motility and improves key neutrophil antimicrobial effector functions.

PTP1B is a non-receptor tyrosine phosphatase that targets a variety of signaling pathways including JAK/STAT, PI3K, and Ras/MAPK ^17^. Thus, PTP1B acts as an intracellular checkpoint to negatively regulate cell responses. PTP1B has been targeted for dendritic cell and CAR T-cell based immunotherapies, in which deletion enhanced cell activation and tumor cytotoxicity ^28,29^. Additionally, specific neutrophil effector functions including migration, phagocytosis, NETosis, and cytokine production have been shown to be affected by PTP1B phosphatase activity ^30-32^. Accordingly, we found that deletion of PTP1B increased PI3K and MAPK signaling and improved iNeutrophil function.

Neutrophils are critical for clearance of fungal infections, including the opportunistic pathogen *A. fumigatus*, which commonly infects immunosuppressed patients ^33^. We found that PTP1B-KO iNeutrophils are significantly better at inhibiting fungal growth over WT cells, due to enhanced recruitment to *A. fumigatus*. PTP1B has been shown to regulate cell motility, but its activating versus inhibitory effects are cell-type and chemokine dependent ^15,16,32^. Here we show that deletion of PTP1B in human iNeutrophils resulted in increased chemotaxis to fMLP. Furthermore, KO iNeutrophils showed enhanced migration in the presence of the fungal pathogen *A. fumigatus*, likely in response to pathogen associated molecular patterns (PAMPs). PTP1B directly targets p38 to inhibit MAPK signaling involved in cell migration towards fMLP ^34^. Indeed, MAPK signaling was increased in stimulated PTP1B-KO iNeutrophils and may promote cell motility towards other stimuli including *A. fumigatus*. Additionally, overexpression of PTP1B results in disorganized distribution of F-actin and focal adhesions ^35^. We found that PTP1B-KO iNeutrophils had increased phosphorylated HS1 and actin polymerization after fMLP stimulation, as well as increased cell polarization in response to *A. fumigatus*. HS1 activates Rac-GTPase signaling and Arp2/3-mediated actin polymerization necessary for cell polarity during chemotaxis ^18,36^. Thus, PTP1B may limit iNeutrophil motility through inhibition of MAPK signaling and actin organization.

During tissue damage and pathogen clearance, neutrophils produce chemotactic factors such as IL-8 and LTB4 to amplify recruitment and promote swarming ^37,38^. We and others have shown that primary neutrophils cluster and produce IL-8 in response to *A. fumigatus* ^13,39,40^, and addition of IL-8 alone can promote primary neutrophil swarming and inhibition of *A. fumigatus* growth ^40^. Here we show that PTP1B-KO iNeutrophils are significantly better at swarming fungal hyphae than WT cells. This is likely due to increased production of IL-8, as this chemokine was elevated with Zymosan stimulation. WT iNeutrophils are capable of swarming but do so infrequently due to limited recruitment to *A. fumigatus*. We found no differences in expression of the LTB4 receptor BLT1R or LTB4 release after fMLP stimulation in WT vs KO cells. Thus, the increased ability of PTP1B-KO iNeutrophils to swarm is likely due to elevated inflammatory signaling initiated upon interaction with fungal hyphae.

Swarming can inhibit fungal growth through release of ROS, NETs and granule proteins such as MPO ^41^. Drug inhibition of PTP1B in murine neutrophils reduces the capacity to NET ^32^. Here, we show that deletion of PTP1B reduces iNeutrophil NET and ROS production in response to PMA stimulation. Although, WT iNeutrophils were more responsive to PMA than KO iNeutrophils, the majority of cells remained inactive in the presence of *A. fumigatus* and were not recruited to hyphae. The differential response of WT iNeutrophils to these two stimuli is likely due to differences in receptor activation and the potency of each stimulus. We predict that PTP1B-null iNeutrophils produce ROS and NETs in response to *A. fumigatus* due to increased cell activation and recruitment to hyphae. Thus, increased cell migration and swarming of PTP1B-KO iNeutrophils likely promotes release of ROS and NETs that are responsible for inhibiting hyphal growth.

Neutrophil phagocytosis is a means for inhibiting fungal growth at the germling stage. PTP1B-KO iNeutrophils readily phagocytosed *A. fumigatus* germlings, correlating to increased uptake of *E. coli* coated beads. Increased actin polymerization in KO cells may promote phagocytosis as remodeling of the actin cytoskeleton is necessary for formation of the phagosome (13). Enhanced TLR signaling may also increase iNeutrophil phagocytosis. While, TLRs are not phagocytic receptors, they can prime neutrophils to promote phagocytosis (32). Accordingly, in response to LPS or Zymosan treatment, we found increased production of key inflammatory mediators, including IL-1β and TNFα. Studies with PTP1B-/- mice have shown increased TLR4 signaling in response to LPS stimulation ^30,42^, supporting the idea that intracellular signaling is increased downstream of PRRs in PTP1B-KO iNeutrophils. Taken together, PTP1B-null iNeutrophils are more responsive to *A. fumigatus* and are capable of inhibiting fungal growth via recruitment and activation of antimicrobial functions including swarming, phagocytosis, NETosis and ROS production.

A caveat of our experiments is the use of the total population of differentiated iNeutrophils. Our flow cytometry analysis of lineage marker expression indicates that iNeutrophils are heterogeneous and vary in their level of neutrophil maturation. In particular, PTP1B-KO iNeutrophils are less mature than WT cells, with lower expression of CD15 and CD16. Immature neutrophils have reduced capacity for NETosis and phagocytosis ^43,44^. Accordingly, we found decreased NET and ROS production in response to PMA stimulation by the total population of PTP1B-KO iNeutrophils. To normalize the maturity level between WT and KO iNeutrophils, we used CD15-positive magnetic bead selection, but saw no improvement in these antimicrobial functions by PTP1B-KO iNeutrophils (data not shown). Thus, the decrease in NETosis and ROS production is likely due to the impact of PTP1B on downstream signaling and not differences in neutrophil maturity. In flow cytometry assays, we can dissect differences in PTP1B-regulated function versus neutrophil maturity by gating on mature neutrophils (CD15+CD16+) only. Through this method, we identified enhanced phagocytosis within mature PTP1B-KO iNeutrophils, but not the bulk population (data not shown). We are currently evaluating methods to generate a more homogenous population of mature PTP1B-KO iNeutrophils.

Here, we identified PTP1B as a negative regulator of iPSC-derived neutrophil cellular activation and effector function. Upon deletion of this phosphatase, we found enhanced intracellular signaling and iNeutrophil function. Our findings suggest that inhibition of PTP1B may be a promising target for further development of iNeutrophil therapies for neutropenic patients with bacterial and fungal infections.

## Data Sharing Statement

All relevant data are included in the manuscript.

## Acknowledgements

Research reported in this publication was supported by the National Institute of Allergy and Infectious Diseases (NIAID) of the National Institutes of Health under Award Number RO1 AI134749-05 and T32AI055397. The content is solely the responsibility of the authors and does not necessarily represent the official views of the NIH. The funders had no role in study design, data collection and analysis, decision to publish, or preparation of the manuscript. The content is solely the responsibility of the authors and does not necessarily represent the official views of the National Institutes of Health.

## Contribution

M.A.G., D.A.B., T.J.S., A.N.P., and H.S.J., performed experiments. M.A.G. D.A.B., T.J.S., A.N.P., J.B, H.Q.D., I.I.S., and A.H. analyzed and interpreted data. M.A.G. and A.H. designed the research and wrote the paper.

