## Supplemental data for "PTP1B phosphatase puts a brake on iPSC-derived neutrophil motility and antimicrobial function"

### Methods

##### Stem Cell Culture and Neutrophil Differentiation

Neutrophils were differentiated from bone marrow derived hiPSCs as previously described (8). Briefly, bone marrow-derived IISH2i-BM9 were obtained from WiCell (Madison, WI). hiPSCs were cultured on Matrigel-coated tissue culture plates in mTeSR-Plus medium (STEMCELL Technologies).

To induce hemogenic endothelium, hiPSCs are transfected with *ETV2* mmRNA in TeSR-E8 media (STEMCELL Technologies) using TransIT reagent and mRNA boost (Mirus Bio). One hour prior to transfection, cells were detached by TrypLE Select (LifeTech). Cells were re-plated onto collagen (2.4ug/ml) coated plates in TeSR-E8 media with 10uM ROCK ROCK inhibitor (ROCKi; Tocris Y-27632). One day following transfection, media was changed to StemLineII media with 20ng/mL VEGF-165 and 10ng/mL FGF to induce differentiation into hemogenic endothelial cells. We refer to this media cocktail as Media A. After two days, the media was changed to differentiate the cells into common myeloid progenitors (CMPs) with StemLineII media supplemented with FGF2 (20 ng/mL; PeproTech), GM-CSF (25 ng/mL; PeproTech), and UM171 (50 nM; Xcess Biosciences). We refer to this media cocktail as Media B. On days 8-10, floating cells were gently harvested and used for terminal neutrophil differentiation. These cells were cultured in StemSpan SFEM II medium (STEMCELL Technologies), supplemented with GlutaMAX 100X (1x; Thermo Fisher Scientific), ExCyte 0.2% (Merck Millipore), human G-CSF (150 ng/mL; Amgen), and Am580 retinoic acid agonist (2.5 µM; Sigma-Aldrich) at $1 \times 10^6$ cells/mL density. We refer to this media cocktail as Media C. After 4 days, fresh Media C was added on the top of the cells. Neutrophils were harvested from the supernatant 8-10 days after plating in Media C.

##### Generation of PTP1B$^{-/-}$ BM9iPSCs

To generate PTP1B knockout BM9 iPSCs, two single guide RNAs were designed using the Synthego CRISPR design tool to target *PTPN1* exon 3: 5’- GATGTAGTTTAATCCGACTA-3’ (sgRNA1) and 5’-TAAAAATGGAAGAAGCCCAA-3’ (sgRNA2). Prior to nucleofection, BM9 iPSCs were detached by TrypLE Select (LifeTech) and singularized by pipetting. 5ug of both sgNRAs and 5ug of Cas9 protein (PNA Bio) were incubated together for 10 minutes, then the cells were nucleofected using the Human Stem Cell Nucleofector Kit 2 (Lonza, #VPH-5022). Cells were plated at 25 cell/cm$^2$ on a Matrigel coated plate in mTeSR-Plus supplemented with 1xCloneR supplement (StemCell #05888). After 2 days, media was changed to mTeSR-Plus only. Individual colonies were picked after 7 days and further expanded. To confirm biallelic mutation in *PTPN1*, genomic DNA was extracted from individual clones, then screened by PCR for the acquisition of a 67 bp deletion using primer1: 5’- TGCATCAGAGAACAGATCCT-3’ and primer2: 5’- CTGGGTAAGAATGTAACTCC-3’. After western blot verification of multiple PTP1B -/- clones, we selected a single clone (#6) for further experimentation.

##### Human Neutrophil Isolation

All blood samples were obtained from healthy donors and were drawn according to the University of Wisconsin-Madison Minimal Risk Research Institutional Review Board-approved protocol (ID: 2017–0032) per the Declaration of Helsinki. Human blood was obtained from volunteering donors with informed written consent through a protocol that was approved by the Internal Review Board of the University of Wisconsin-Madison. Neutrophils were isolated using MACSxpress negative antibody selection kit and purified with the MACSxpress erythrocyte depletion kit (Miltenyi Biotec, Inc., Auburn, CA), following manufacturer’s instructions. Isolated neutrophils were resuspended in modified HBSS

(+0.1% HSA +10mM HEPES) and utilized for various experiments. Incubations involving neutrophils were performed at 37°C with 5% $CO_2$.

**Cytospins**

To confirm neutrophil morphology, 90,000 cells were spun onto a glass slide for 5 minutes at 1200rpm using a Shandon Cytospin 3 centrifuge (ThermoFisher). Cytospin slides were then stained with Differential Quick III Stain Kit (Electron Microscopy Sciences cat#26096-25) following manufacturer's instructions. Slides were imaged at 20x on an Olympus IX70 microscope and evaluated for neutrophils by the presence of hypersegmented nuclei.

**Flow Cytometry**

Flow cytometry analysis was used to evaluate expression of cell lineage markers on iPSC-derived neutrophils. Neutrophils were stained in PBS+ 1%HSA media +Brilliant Buffer (Thermo Fisher #00-4409-42) and Human TruStain FcX Fc Receptor Blocking Solution (Biolegend #422302), then fixed with 2% PFA. Data acquisition was performed on an Aurora Cytometer (CytekBio). Antibodies used in this study can be found in **Supplementary Table 1.** Forward and side-scatter parameters identified single cells and live cells were identified with Ghost Dye Red 780 or Zombie NIR dye. Myeloid cells were identified by CD11b+ expression and mature neutrophils were identified by CD15+ or CD15+CD16+ expression. Monocytes were identified as CD14+ cells. Data were analyzed using FLowJo Software (v10.8.1).

*Flow cytometry analysis in R using Cytofworkflow pipeline*

Flow cytometry (.fcs) files were read into FlowJo (v10.8.1) after spectral unmixing. Initial gates using forward and side-scatter parameters isolated single cells, and a viability dye was used to exclude dead or dying cells. For each sample, live, single cells were exported, selecting the 'all compensated parameters' option to generate new .fcs files in FlowJo for further analysis using R (v4.0.3) with *prepData* function from Cytofworkflow (v1.14.0) pipeline (9). Data transformation was calculated for all fluorescent parameters using *arcsinh* normalization with a cofactor of 3000. The distribution of marker expression for transformed data was plotted using a smoothed density function using ggplot2 to compare WT and KO samples.

**Presto Blue Viability**

Primary human neutrophils and iPSC-derived neutrophils were resuspended in RPMI media supplemented with 10%FBS. 100k cells were plated into each well of a black clear-bottom 96-well plate. Separate plates were prepared for each timepoint (days 0, 1, 3 and 5). On day 0, cells were allowed to rest for at least 1 hour prior to addition of Presto Blue HS (Thermo Fisher #P50200). For each timepoint, Presto Blue HS was added and incubated for 30 minutes at 37°C before reading fluorescence at 560/590 nm in a microplate reader (Synergy H1, Bio-Tek Instruments). Background fluorescence of media only wells was subtracted from each sample and the fold change was calculated compared to day 0, respective to each cell line.

**Western Blot Cell Signaling**

Western blotting was conducted to quantify PTP1B protein expression and evaluate phospho-signaling in our iPSC-derived neutrophils. Cells were stimulated with 1uM fMLP for 3 minutes. Cell pellets were collected in lysis buffer (25 mM HEPES, pH 7.5, 150mM NaCl2, 1% Nonidet P-40, 10mM MgCl2, 1mM EDTA, 10% glycerol supplemented with 1g/ml pepstatin A, 2g/ml aprotinin, 1 g/ml leupeptin, and 200nM phenylmethanesulfonyl fluoride, 1mM sodium orthovanadate, 1% protease inhibitor mixture 2 (Sigma #p5726)). Cells were incubated on ice for 10 minutes and then sonicated with 20% amplitude for

3x 5 sec. Cells were then clarified by centrifugation at 15,000 ×g, 4°C for 15 min. Protein concentrations were determined using the Pierce BCA Protein Assay (Thermo Scientific; 23225) and samples stored at −80°C. Immunoblotting of cell lysates was performed and blots were imaged with an infrared imaging system (Odyssey; LI-COR Biosciences). Primary and secondary antibodies used can be found in **Supplementary Table 2**.

**Chemotaxis**

Chemotaxis was assessed using a microfluidic device as described previously (10). In brief, polydimethylsiloxane (PDMS) devices were plasma treated and adhered to glass coverslips. Devices were coated with 10 µg/mL fibrinogen (Sigma) in PBS for 30 min at 37°C, 5% $CO_2$. The devices were blocked with 2% BSA-PBS for 30 min at 37°C, 5% $CO_2$, and then washed twice with modified HBSS (+0.1% HSA +10mM HEPES) (mHBSS). Cells were stained with Calcein AM (Molecular Probes) in PBS for 10 min at room temperature followed by resuspension in modified HBSS (+0.1% HSA +10mM Hepes). Cells were seeded at $5 \times 10^6$/mL to allow adherence for 30 min before addition of chemoattractant. Then, 3uL of 1 µM fMLP (Sigma) chemoattractant was loaded into the input port of the microfluidic device. Cells were imaged every 30 seconds for 45–90 min on a Nikon Eclipse TE300 inverted fluorescent microscope with a 10× objective and an automated stage using MetaMorph software (Molecular Devices). Automated cell tracking analysis was done using JEX software (11) to calculate chemotactic index and velocity and to generate rose plots.

**Immunofluorescent imaging**

iNeutrophils were stimulated and prepared for immunofluorescent imaging as previously described (12). Acid-washed 22-mm glass circle coverslips were coated with 10 µg/ml fibronectin for at least 1 h at 37°C and then blocked for 30 min with 2% BSA-PBS. Cells ($3 \times 10^5$) in 500 µl 0.5%HSA-RPMI were seeded per coverslip in a 24-well plate and allowed to rest for 30 min. fMLP was added for a final concentration of 100 nM fMLP and cells were allowed to adhere for 30 minutes at 37°C. Media was aspirated and fixation was performed as follows: 1 ml of 37°C preheated 4% paraformaldehyde in PEM buffer (80mM PIPTES, pH6.8; 5mM EGTA, pH 8.0; 2mM $MgCl_2$) for 15 min at RT. Fixative was aspirated and 0.25% Triton X-100 in PEM buffer was added for 10 min at 37°C. Coverslips were washed three times with PBS, blocked in 5%BSA-PBS for 60 min at RT, washed, and incubated in Rhodamine-phalloidin (Thermo Fisher Cat#R415) overnight at 4°C. Coverslips were washed, counterstained with Hoechst 33342 (Thermo Fisher H3570; 1:500) for 5–10 min, washed with dd$H_2O$, and mounted on Rite-On Frosted Slides (Fisher Scientific; 3050-002) with ProLong Gold Antifade Mountant (Invitrogen; P36930). Slides were imaged on an upright Zeiss LSM 800 Laser Scanning Confocal Microscope equipped with a motorized stage and Airyscan module. Images were acquired with a 60× oil, NA 1.40 objective using the Airyscan acquisition mode. Images were processed in ZenBlue software (Zeiss) using the Airyscan processing method and maximum intensity projections generated for further analysis. Integrated density of the actin channel was quantified using ImageJ.

**Phagocytosis**

Phagocytosis was quantified using pHrodo™ Green *E. coli* BioParticles™ (Invitrogen #P35366) following manufacturer's instructions. Briefly, the *E. coli* BioParticles were opsonized with 30% pooled human serum (MP Biomedicals #MP092930149) for 30 minutes at 37°C then washed 3 times in PBS. 1 million cells were resuspended in 80uL of Media C and 20uL of opsonized beads were added at 100:1 MOI. Cells and beads were incubated for 1 hour at 37°C, then stopped by addition of ice cold PBS. While keeping tubes on ice, cells were stained with neutrophil lineage markers (**Supplementary Table 1**) then fixed with 2% PFA before flow cytometry analysis on the Aurora Cytometer (CytekBio).

**Intracellular ROS**

Intracellular peroxynitrite production by iPSC-differentiated neutrophils was quantified using the peroxynitrite indicator DHR123 (Invitrogen # D23806). A black wall clear-bottom 96-well plate was pre-coated with 10ug/ml fibronectin. 100,000 cells in Phenol Red Free-RPMI containing 2%FBS and 5ug/mL DHR123 reagent were plated into each well. Cells were allowed to rest for 30 minutes at 37°C. PMA at a final concentration of 50ng/mL was added to appropriate wells. Wells were plated in quadruplicate to account for technical error. Reads were taken every 15 minutes for 2 hours at 485/535nm using the Victor3V microplate reader (PerkinElmer). Background signal of unstimulated cells was subtracted from the corresponding PMA stimulated cell line and plotted over time. Samples were normalized to the percent of maximal ROS production by WT iNeutrophils to account for daily fluctuations in RFUs.

**NETosis Assay**

A black walled clear-bottom 96-well plate was pre-coated with 10ug/mL Fibronectin. 200,000 cells in 100uL Phenol Red Free RPMI + 2%FBS were plated into each well. Cells were allowed to rest for 30 minutes at 37°C. Final concentration of 100ng/mL PMA was added to appropriate wells and the plate was incubated for 4 hours at 37°C. Sytox Green (Invitrogen # S7020) was added at 375nM final concentration. The microplate was incubated for 10 minutes and an endpoint reading was taken using a Victor3V microplate reader (PerkinElmer) to quantify extracellular DNA by fluorescence (500/528nm). Background signal of Sytox Green unstimulated cells was subtracted from the corresponding PMA stimulated cell line. Fold change of fluorescence was calculated compared to WT.

**CD15 Positive Selection**

hiPSC-derived neutrophils were positively selected for mature neutrophils using surface expression of CD15. CD15 microbeads (Miltenyi Cat# 130-046-601) were incubated with neutrophils following the manufacturer's protocol and then positively selected using LD columns (Miltenyi Cat# 130-042-901). Cells were allowed to rest overnight in Media C with added 10%FBS before use in further experiments.

**Real Time qPCR**

iPSC-differentiated neutrophils were stimulated with LPS to evaluate expression of inflammatory cytokines. A 6 well TC treated plate was pre-coated with 10ug/ml fibronectin. Approximately 6 million cells resuspended in 2%FBS-RPMI were plated per well. Cells were allowed to rest for 30min at 37˚C before stimulation. Final concentration of 200ng/mL *E. coli* LPS (Sigma # L2755) was added to appropriate wells and the plate was incubated for 2 hours at 37˚C. Floating cells were collected and spun down at 300xg. To collect RNA, 1ml of Trizol (Invitrogen) was added to adherent cells and then combined with the pelleted floating cell sample. Samples were pipetted to mix and then stored at -80°C until RNA extraction. RNA was isolated by Trizol extraction following manufacturer's instructions (Invitrogen #15596026). Collected RNA was stored at -80°C until cDNA preparation. cDNA was synthesized using oligo-dT primers and the Superscript III First Strand Synthesis kit (Invitrogen #18080-051) following manufacturer's instructions. cDNA was used as the template for quantitative PCR (qPCR) using FastStart Essential Green DNA Master (Roche) and a LightCycler96 (Roche). Data were normalized to *ef1a* within each sample using the ΔΔCq method (13). Fold-change represents the change in cytokine expression over the unstimulated WT sample. All qPCR primers are listed in **Supplementary Table 3**.

**Inflammatory Cytokine and LTB4 ELISA**

iNeutrophil secretion of inflammatory cytokines IL-8, IL-1beta, IL-6 and TNFa protein was quantified by ELISA. A 12-well TC treated plate was coated with 10ug/mL fibronectin. iNeutrophils were resuspended in 2%FBS supplemented RPMI and 1.25-1.5 million cells were plated per well. Cells were allowed to rest for 30min at 37°C before stimulation with 200ng/mL *E. coli* LPS or 10ug/mL Zymosan for 4 hours at 37°C.

Wells were harvested and the media was spun down at 300xg to pellet the cells. Supernatants were collected, aliquoted and frozen at -80°C until use. Cytokines were quantified with the Human IL-8/CXCL8 DuoSet ELISA (Bio-Techne DY208), Human IL-6 DuoSet ELISA (Bio-Techne DY206), Human TNF-alpha DuoSet ELISA (Bio-Techne DY210), and Human IL-1 beta/IL-1F2 DuoSet ELISA (Bio-Techne DY201) following manufacturer's instructions. Samples were diluted to fit in the range of the standard curve and normalized to the input cell number.

LTB4 release was quantified following stimulation with fMLP. A 24-well TC treated plate was coated with 10ug/mL fibronectin. iNeutrophils were resuspended in 0.5%HSA supplemented RPMI and 1 million cells were plated per well. Cells were allowed to rest for 30min at 37°C before stimulation with 1uM fMLP for 5 minutes or 30 minutes. Wells were harvested and the media was spun down at 300xg to pellet the cells. Supernatants were collected, aliquoted and frozen at -80°C until use. LTB4 release was quantified with the LTB4 Parameter Assay Kit (Bio-Techne KGE006B) following manufacturer's instructions.

**Fungal co-culture imaging**

*Aspergillus fumigatus* (CEA10) was grown as previously described (14). Briefly, *A. fumigatus* (CEA10) was grown on glucose minimal medium (GMM) plates at 37°C in the dark to promote asexual conidiation. Aspergillus was plated at $1x10^6$ conidia/10cm plate for 3-4 days. Conidia were harvested in 0.01% Tween water by scraping with an L-spreader and then passed through sterile Miracloth into a 50 ml conical tube. The spore suspension was centrifuged at 900 x*g* for 10 min at room temperature and re-suspended in 50 ml 1× PBS. The spore suspension was then vacuum filtrated using a Buchner filter funnel with a glass disc containing 10–15 µm diameter pores. The filtered suspension was centrifuged at 900 x*g* for 10 min and re-suspended in 1 ml 1× PBS. Conidia were counted using a hemacytometer and the concentration was adjusted to $1.5\times10^8$ spores/ml. Conidial stocks were stored at 4°C and used up to 1 month after harvesting.

Live imaging was conducted to visualize iPSC-derived neutrophil interactions with fungal hyphae. *A. fumigatus* 2x $10^3$ spores/well were plated in 100uL GMM media in a black 24-well plate coated with fibronectin (Corning). The plate was incubated at 37°C for 8 hours, or until germling stage. Spore germination was confirmed by microscopy prior to adding neutrophils. iPSC-derived neutrophils were resuspended in RPMI + 2%FBS at $6x10^5$ cells/mL. GMM media was removed from the wells and replaced with 100uL of neutrophil suspension to yield a neutrophil to spore ratio of 150:1. Neutrophil-fungal interactions were imaged every 3 minutes on an inverted fluorescent microscope (Nikon Eclipse TE300) with a 20× objective and an automated stage (Ludl Electronic Products) with a Prime BSI Express camera (Teledyne Photometrics). Environmental controls were set to 37°C with 5%C $CO_2$. Videos were compiled using ImageJ software.

*Image Analysis*

iNeutrophil circularity was quantified after 0 and 4 hours of co-incubation with *A. fumigatus*. Individual cells were outlined using the Polygon selection tool in ImageJ and circularity ($4\pi$*area/perimeter$^2$) was calculated and is reported in Figure 5C. The hyphal length for individual germlings was measured after 0 and 4 hours of co-incubation with iNeutrophils. Germlings and hyphae were traced using the segmented line feature in ImageJ and the cumulative lengths of individual germlings are reported in Figure 5G. The percent of germlings with clustered iNeutrophils and cluster size was quantified after 0, 1, 2, and 4 hours of co-incubation with *A. fumigatus*. A cluster was identified as a tightly formed group of at least 5 iNeutrophils attached to a germling or hyphal branch and outlined using the Polygon selection tool (ImageJ). Cluster size and percent of clustered germlings are reported in Figure 5E,F.

**Statistical Analysis**

All experiments and statistical analyses represent at least three independent replicates (N). The replicate number of cells or germlings (n) for experiments in Figure 2F and 5C, G is indicated in the figure legend. Analysis of chemotactic index and cell velocity was performed using unpaired Student's t-test. Analysis of receptor expression by flow cytometry, phagocytosis, qPCR gene expression and ELISA was performed using paired Student's t-test. Analysis of Presto Blue viability, intracellular reactive oxygen species (ROS) production, and percent of clustered germlings was performed using simple linear regression and the $p$ value was calculated by comparing the slope of each line. Analysis of cluster size over time was performed by calculating the area under the curve (AUC) for each replicate and then determining the $p$ value using a paired Student's t-test. Analysis of NETosis and phospho-signaling was performed using One Sample t-test. The above analyses were conducted using GraphPad Prism (v9). Integrated Density of F actin, circularity, and hyphal length analyses represent least-squared adjusted means±standard error of the mean (LSmeans±s.e.m.) and were compared using ANOVA with Tukey's multiple comparisons (RStudio). All graphical representations of data were created in GraphPad Prism (v9) and figures were ultimately assembled using Adobe Illustrator (Adobe version 23.0.6).

#### Supplementary Tables

**Supplementary Table 1. Flow Cytometry antibodies and dyes used in this study**

| Target | Fluor | Clone | Cat # | Vendor |
|---|---|---|---|---|
| CD14 | Spark Blue 550 | 63D3 | 367148 | Biolegend |
| CD16 | AF700 | 3G8 | 302026 | Biolegend |
| CD66b | PE/Fire640 | 6/40c | 392918 | Biolegend |
| CD182 (CXCR1) | BV605 | 5A12 | 743421 | FisherScientific |
| CD182 (CXCR2) | BV605 | 5A12 | 744197 | FisherScientific |
| CXCR4 | APC | 12G5 | 306510 | Biolegend |
| CD10 | PE | HI10a | 312203 | Biolegend |
| CD11b | PECy7 | ICRF44 | 301321 | Biolegend |
| CD15 | APC-Fire 810 | W6D3 | 323058 | Biolegend |
| Zombie NIR | 746 | NA | 423105 | FisherScientific |
| CD11b | PE | M1/70 | 101208 | Biolegend |
| CD15 | APC | W6D3 | 323008 | Biolegend |
| CD16 | BV711 | 3G8 | 563127 | BD Biosciences |
| CD14 | BUV805 | M5E2 | 612902 | BD Biosciences |
| Ghost Dye Red | 780 | NA | 13-0865-T100 | Tonbio Biosciences |
| BLT1R | BUV805 | 14F11 | 749044 | BD Biosciences |
| Clec7a (Dectin-1) | APC | 15E2 | 355405 | Biolegend |
| TLR2 | FITC | W15145C | 392307 | Biolegend |
| TLR4 | PE | HTA125 | 312805 | Biolegend |
| Phrodo Green AM Intracellular pH Indicator | 509/533 | NA | P35373 | Thermo Fisher |
| pHrodo Green *E. coli* Bioparticles | 509/533 | NA | P35366 | Thermo Fisher |

| | | | | |
|---|---|---|---|---|
| Brilliant Buffer | NA | NA | 00-4409-42 | Thermo Fisher |
| Human TruStain FcX Fc Receptor Blocking Solution | NA | NA | 422302 | Biolegend |
| Ultracomp ebeads | NA | NA | 501129040 | Thermo Fisher |

**Supplementary Table 2. Immunoblot antibodies utilized in this study**

| **Target** | **Vendor** | **Catolog No.** |
|---|---|---|
| Anti-Human PTP1B | BD Transduction Laboratories | 610140 |
| Rabbit Anti-Human AKT | Cell Signaling | 9272S |
| Mouse Anti-Human phopsho AKT | Cell Signaling | 4051 |
| Rabbit Anti-Human HS1 (D83A8) | Cell Signaling | 3890S |
| Rabbit Anti-Human phospho HS1 Y397 | Cell Signaling | 4507S |
| Rabbit Anti-human ERK | Thermo Fisher | 44-654G |
| Mouse Anti-Human phospho ERK | Abcam | ab50011 |
| Goat anti-mouse IgG (H&L) Antibody Dylight 800 Conjugated | Rockland Immunochemicals | 610-140-002-0.5 |
| Goat anti-Rabbit IgG (H&L) Antibody Cross-Adsorbed Secondary Antibody, 680 Conjugated | Invitrogen | A-21076 |

**Supplementary Table 3. qRT-PCR primers used in this study**

| Gene | Sequence (5’ → 3’) | Reference |
|---|---|---|
| EF1alpha | F- TGGTATTGGTACTGTTCCTG | KiCqStart SYBR Green Primers |
| | R- CTTCACTCAAAGCTTCATGG | |
| IL1beta | F- CTAAACAGATGAAGTGCTCC | KiCqStart SYBR Green Primers |
| | R- GGTCATTCTCCTGGAAGG | |
| IL6 | F- TCTCCACAAGCGCCTTCG | PrimerDB |
| | R- CTCAGGGCTGAGATGCCG | |
| IL8 | F- TGTAAACATGACTTCCAAGC | KiCqStart SYBR Green Primers |
| | R- AAAACTGCACCTTCACAC | |
| TNFa | F- GACAAGCCTGTAGCCCATGT | Self-designed |
| | R- TCTCAGCTCCACGCCATT | |

#### Supplementary Figures

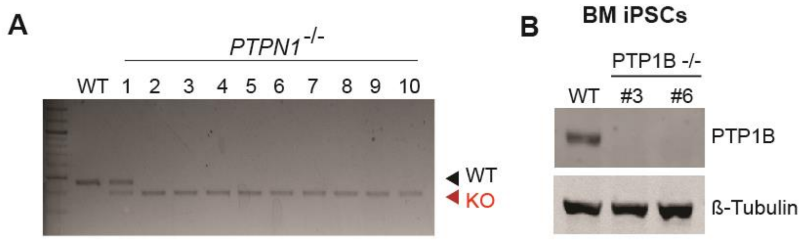


**Supplementary Figure 1. CRISPR/Cas9 mediated deletion of PTP1B.**

(A) Genomic PCR confirmation of a biallelic 67bp deletion in the *PTPN1* gene in 10 clonal lines. (B) Representative western blot showing confirmation of loss of PTP1B protein in bone marrow (BM)-derived iPSC-stem cells.

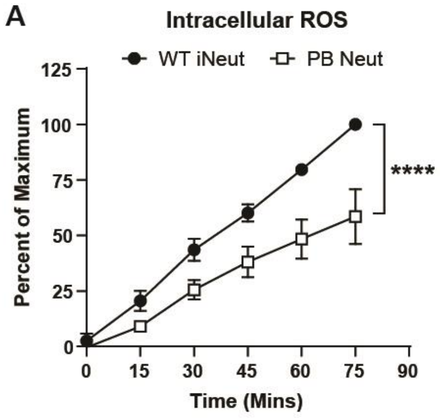


**Supplementary Figure 2. iNeutrophils produce more ROS than peripheral blood neutrophils.**

(A) Quantification of iNeutrophil and human peripheral blood (PB) neutrophil intracellular ROS production over time using DHR123 peroxynitrite indicator following stimulation with 50ng/mL PMA. Experiments were conducted three times. Means ± SEM are shown. *p* values were calculated by simple linear regression. ****, $p < 0.0001$.

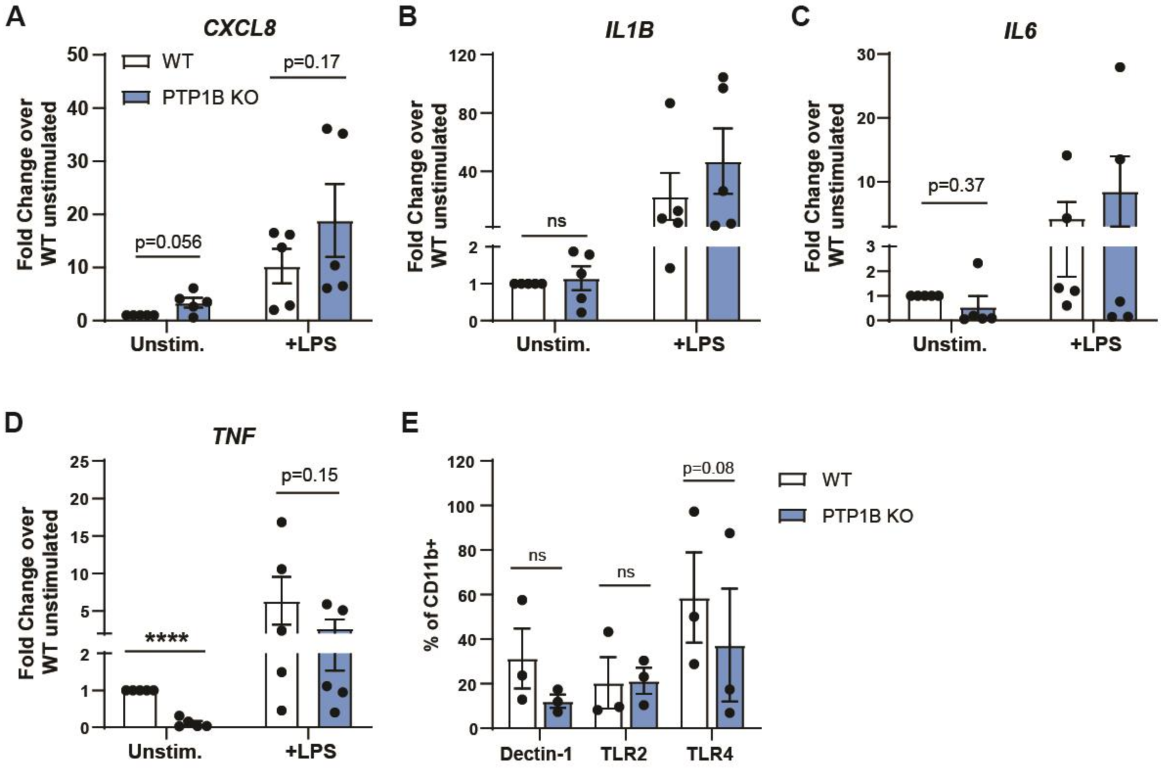


**Supplementary Figure 3. Inflammatory cytokine transcription and Pathogen Recognition Receptor expression on iNeutrophils.**

(A-D) qRT-PCR of inflammatory cytokines produced by unstimulated iNeutrophils or after stimulation with 200ng/mL LPS for 2 hours. qRT-PCR samples were normalized to the WT unstimulated control. (E) Flow cytometry staining for pathogen recognition receptors. Percent was quantified of CD11b+. Experiments were conducted at least three times, or as indicated on the plot. Dots represent independent replicates. Means ± SEM are shown. *p* values were calculated by paired Student's t-test (A-H). ****, $p < 0.0001$.

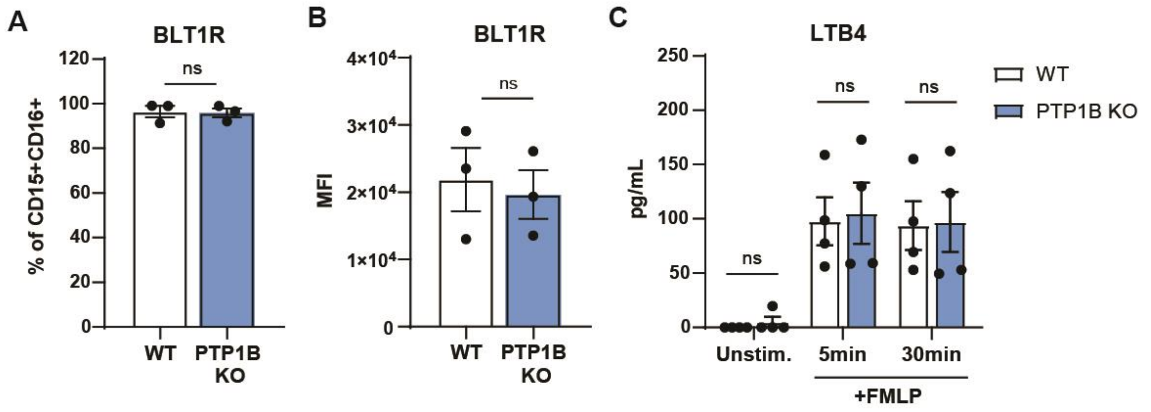


**Supplementary Figure 4. LTB4 receptor expression and release from iNeutrophils.**

(A) Flow cytometry staining for the LTB4 receptor BLT1R. Percent was quantified of CD15+CD16+ iNeutrophils. (B) Quantification of mean Fluorescence Intensity (MFI) for BLT1R expression on CD15+CD16+ iNeutrophils. (C) LTB4 release following 1uM fMLP stimulation for 5 or 30 minutes quantified by ELISA. Experiments were conducted at least three times, or as indicated on the plot. Dots represent individual replicates. Means ± SEM are shown. *p* values calculated by paired Student's t-test (A-C).

#### Videos

**Video 1. PTP1B-KO iNeutrophils migrate towards and cluster on *A. fumigatus* germlings and hyphae**

Representative movie of PTP1B-KO iNeutrophils engaging with *A. fumigatus* germlings and hyphae. Note that at 6 minutes, a germling is phagocytosed. At 9 minutes and 48 minutes, KO iNeutrophils swarm and cluster hyphae that are too large to phagocytose. Germlings are indicated with black arrows. Phagocytosed germling is indicated with a white arrow. Asterisks indicate the initiation of an iNeutrophil cluster. Scale bar is 50µm. Images were taken every 3 minutes. Movie is displayed as 2 frames/second.

**Video 2. WT iNeutrophils show minimal response to *A. fumigatus* germlings**

Representative movie of WT iNeutrophil interaction with *A. fumigatus* germlings. WT iNeutrophils are minimally responsive to *A. fumigatus* with only a fraction of cells migrating towards and interacting with germlings. Note only the center germling is phagocytosed, indicated with a white arrow. Even after 2 hours, clustering is not initiated for any germlings. Germlings are indicated with black arrows. Scale bar is 50µm. Images were taken every 3 minutes. Movie is displayed as 2 frames/second.

**Video 3. WT iNeutrophils show minimal response to *A. fumigatus* hyphae**

Representative movie of WT iNeutrophil interaction with *A. fumigatus* hyphae. WT iNeutrophils are minimally responsive to *A. fumigatus* with only a fraction of cells migrating and clustering on hyphae. The top hypha shows iNeutrophil recruitment after two hours of co-incubation, resulting in cluster formation (indicated with an asterisk). The bottom hypha shows no recruitment and nearby iNeutrophils remain inactive even after 2 hours. Scale bar is 50µm. Images were taken every 3 minutes. Movie is displayed as 1 frame/second.

**Video 4. PTP1B-KO iNeutrophils rapidly migrate towards and cluster on *A. fumigatus* germlings.**

Representative movie of PTP1B-KO iNeutrophils engaging with an *A. fumigatus* germling. Note that the germling is rapidly phagocytosed and results in cluster formation. Germling is indicated with a black arrow. Blue arrow indicates a cell migrating towards the germling. White arrow indicates a phagocytosed germling. Asterisk indicates the formation of an iNeutrophil cluster. Scale bar is 50µm. Images were taken every 3 minutes. Movie is displayed as 1 frame/second.

**Video 5. PTP1B-KO iNeutrophils rapidly migrate towards and cluster on *A. fumigatus* germlings.**

Representative movie of PTP1B-KO iNeutrophils engaging with an *A. fumigatus* germling. Note that the germling is rapidly phagocytosed and results in cluster formation. Germling is indicated with a black arrow. Blue arrow indicates a cell migrating towards the germling. White arrow indicates a phagocytosed germling. Asterisk indicates the formation of an iNeutrophil cluster. Scale bar is 50µm. Images were taken every 3 minutes. Movie is displayed as 1 frame/second.
